# Single-session measures of quadriceps neuromuscular function are reliable in healthy females and unaffected by age

**DOI:** 10.1101/2023.06.22.545905

**Authors:** Steven J. O’Bryan, Danielle Hiam, Séverine Lamon

## Abstract

**Objective:** The inter-session reliability of a wide range of measures used to characterize the aging neuromuscular system is unknown, particularly in females. The aim of this study was to determine the inter-session reliability of quadriceps neuromuscular function assessed via maximal voluntary and evoked force and electromyography responses in healthy young and older females.

**Methods:** Twenty-six females aged 19 – 74 years completed two identical testing sessions 9 ± 7 days apart. Quadriceps neuromuscular function measurements included isometric maximal voluntary force (MVC), high and low frequency twitch force, voluntary and evoked electromyography (EMG) in superficial quadriceps (RMS, M-wave and H-reflex), and maximal torque (T0), velocity (V0) and power (P_MAX_) derived from torque-velocity and power-velocity relationships. Intra-class correlation coefficients (ICC), coefficients of variation (CoV) and Bland-Altman plots were used to assess inter-session reliability. The effect of participant age on inter-session reliability was assessed by linear regression.

**Results:** Excellent reliability (ICC > 0.8) was shown for all voluntary and evoked mechanical outcomes and systematic bias was essentially absent. Similarly, all *vastus lateralis* EMG outcomes showed excellent reliability (ICC > 0.8) with CoVs < 12%, which were better than *vastus medialis* and *rectus femoris* outcomes. Participant age was not associated with inter-session reliability (P > 0.05).

**Conclusion:** Excellent reliability of voluntary and evoked force and *vastus lateralis* EMG outcomes measured in healthy females can be attained in one testing session, irrespective of age, increasing feasibility for future research. The random error should however be considered when quantifying age-related differences and/or adaptation to exercise in female neuromuscular function.

**New and Noteworthy:** The test-retest reliability of a diverse range of measures used to quantify neuromuscular function were assessed in younger and older females for the first time. We show that reliable measures of maximal voluntary and evoked quadriceps force and electromyography outcomes can be obtained in one testing session, irrespective of participant age. Thus, neuromuscular function can be accurately assessed across the female lifespan with minimal inconvenience imposed on participants, increasing feasibility for future research.

## 1. Introduction

The degenerative effects of the aging neuromuscular system and the effectiveness of interventional strategies aimed at improving motor function are grounded in cross-sectional or longitudinal observations. For example, some studies demonstrate an annual reduction in strength of ∼1-4% past the age of 65 years (1, 2), whereas others show a ∼15-40% reduction in muscle shortening velocity and a ∼10–30% reduction in power in older compared to younger adults (3, 4). Yet, no studies have concomitantly evaluated the inter-session reliability of a wide range of valuable measures used to assess neuromuscular function at different stages of the lifespan. Moreover, although more research specific to female neurophysiology is emerging, test-retest reliability of neuromuscular function has not been established in females despite unique physiological profiles known to influence the neuromuscular system (5-7).

Neuromuscular function is characterized by the translation of synaptic inputs received by motoneurons into the generation of external forces necessary for human movement (8). At a general level, neuromuscular function may assessed by quantifying maximal force/torque generated during isometric (7, 9), isokinetic (10), or isoinertial (11) voluntary contractions. In each instance, reported inter-session reliability was high in younger and older males and females (intra-class correlation coefficient > 0.88) (12-21), although isoinertial contractions may be less reliable (22). Despite high reliability of such measures of muscle strength, these tests are limited in their capacity to concurrently evaluate maximal shortening velocity and power production, which is arguably closer related to motor function than maximal strength (23, 24). Thus, others have evaluated age-related deficits in neuromuscular function by modeling torque-velocity (T-V) and power-velocity (P-V) relationships derived from a series of maximal dynamic contractions (single or multi-joint) performed at different isoinertial loads or isokinetic velocities (25-29). However, the reliability of T-V and P-V outcomes from a large functional muscle such as the quadriceps has not been established. Quadriceps function is strongly related to lower-limb mobility and power production across the lifespan (27-31). Coupling T-V and P-V relationships of quadriceps with techniques such as electrical nerve stimulation and surface electromyography (EMG) provides deeper knowledge regarding the neurophysiological properties that govern maximal force, velocity and power, beyond what can be obtained from maximal voluntary contractions (32).

A superimposed stimulus applied to the nerve during maximal voluntary contraction can evaluate the capacity of cortical and spinal motoneurons to activate muscle fibers and achieve maximal voluntary force (i.e. voluntary activation) (33), whereas high and low frequency stimulation at rest can evaluate mechanisms which influence neuromuscular transmission and excitation-contraction coupling (32, 34, 35). Moreover, EMG recordings of compound muscle action potentials (i.e. M-wave) and Ia afferent reflexive responses (i.e. H-reflex) provide details on the capacity and velocity of action potential propagation (32, 36) and muscle spindle excitation of alpha motoneurons (37). Despite the considerable value of electrical stimulation in assessing neuromuscular function in older populations (34, 38, 39), the inter-session reliability of its outcomes have primarily been investigated in younger males (15-21) and once in older males and females with knee osteoarthritis (40). Age and sex both have influence on the neurophysiological and morphological features of the superficial quadriceps (5, 41, 42), indicating that inter-muscular differences may exist in the inter-session reliability of EMG outcomes and warranting more specific investigations.

The aim of this study was to determine the test re-test reliability of quadriceps neuromuscular function in healthy females between 18 – 80 years of age. Maximal isometric contractions, low and high frequency doublets, force-velocity and power-velocity relationships during dynamic contractions, and voluntary and evoked surface electromyography responses (RMS, M-wave, H-reflex) were assessed during two identical testing-sessions. Additionally, inter-muscular differences in the reliability of EMG outcomes between the superficial quadriceps was evaluated.

## 2. Materials and Methods

### 2.1 Participants

Sample size was calculated with the method presented by Borg et al. (43) and determined from expected intraclass correlation coefficients (ICC) previously reported for similar outcomes and population sample (12, 13). In this way, a required sample size of twenty-three was calculated with an expected ICC of 0.9, minimum ICC of 0.7, and statistical power of 80%. An additional 10% were recruited to account for participant drop out or missing data points. Twenty-six healthy females (sex chromosome complement XX) aged 19 – 74 years (mean ± standard deviation: age = 43 ± 18 years, height = 164 ± 5.7 cm, weight = 63.5 ±10.8 kg, BMI = 23.5 ± 3.9 kg.m^2^) volunteered to participate in this study. Two participants did not engage in physical exercise, eight performed light intensity exercises (e.g. pilates, golf, walking), nine were moderately trained (e.g. resistance exercise, jogging) and seven regularly performed vigorous exercise (competitive hockey, high-intensity running, cross-training). Seventeen participants were pre-menopausal, with ten of those with an implanted intrauterine device or taking oral contraceptives. The remaining ten participants were post-menopausal with one participant on hormone replacement therapy. Prior to participation, a medical history and risk-factor assessment questionnaire was completed to confirm eligibility. The exclusion criteria included pregnancy, cancer, implanted medical devices, BMI > 35kg.m^2^, or any diseases of the central nervous system, musculoskeletal system, cardiorespiratory system or metabolic system. Written informed consent was obtained from each participant prior to commencement of the study. All testing procedures were approved by the Deakin University Human Research Ethics Committee (DUHREC 2021-307).

### 2.2 Experimental Protocol

Participants visited the laboratory on two separate occasions at the same time of day, with a duration of 9 ± 7 (mean ± standard deviation) days separating each visit. Participants were instructed to avoid strenuous physical activity in the 48 hours preceding each visit, and to avoid caffeine on the day of testing.

All experimental testing was conducted on an isokinetic dynamometer (Universal Pro Single Chair model 850-230, Biodex Medical Systems, United States). Participants sat upright in the dynamometer chair with straps across the thorax and pelvis, with hip angle set at 85° flexion. The axis of rotation of the dynamometer was aligned with the axis of rotation of the dominant knee (estimated by palpation of the lateral femoral epicondyle), with the distal aspect of the dynamometer attachment fixed to the leg via Velcro strap ∼2cm proximal to the lateral malleoli of the fibula. Participants were instructed to cross their arms across their chest during all testing.

Following a standardized warm-up consisting of a series of incremental submaximal and maximal isometric knee extensions (one 4s contraction at 20%, 40%, 60% and 80% perceived effort and up to three at 100% perceived effort), maximal voluntary and evoked mechanical and electromyography responses were measured in quadriceps with three experimental protocols separated by 15-min of rest (**Figure 1**). First, participants completed a ⁓4s maximal isometric voluntary contraction (MVC) at 75° knee flexion with an electrically evoked doublet (100Hz) applied to the femoral nerve at the plateau in voluntary force, followed ∼2s after by three resting evoked responses (100Hz, 10Hz and 1Hz ∼1.5s apart) (i.e. twitch interpolation). This procedure was repeated three times with two minutes rest separating each set. Second, Hoffmann (H) reflexes were elicited in quadriceps by applying a 1Hz electrical stimulus to the femoral nerve at progressively increasing intensity during 50 brief (2-3s) submaximal isometric contractions performed at 5% MVC (∼10s rest separated contractions with inter-stimulus duration = 13.6 ± 0.9s). Participants were asked to keep their head straight with eyes on the torque feedback screen, place their arms across the chest, avoid any unnecessary limb movements and remain quiet. Finally, torque-velocity and power-velocity relationships of quadriceps were derived from maximal voluntary contractions performed at five different isotonic loads (∼0% MVC, 15% MVC, 30% MVC, 45% MVC and 50% MVC) and one isokinetic speed (1.047 rad/s). Participants were instructed to extend their knee as hard and fast as possible from 110° - 10° knee flexion (0° = full extension), with two efforts performed for each load/speed (⁓3 s apart) and 3 min rest between different load/speed. Strong verbal encouragement was provided for all maximal efforts.

**Figure 1.**
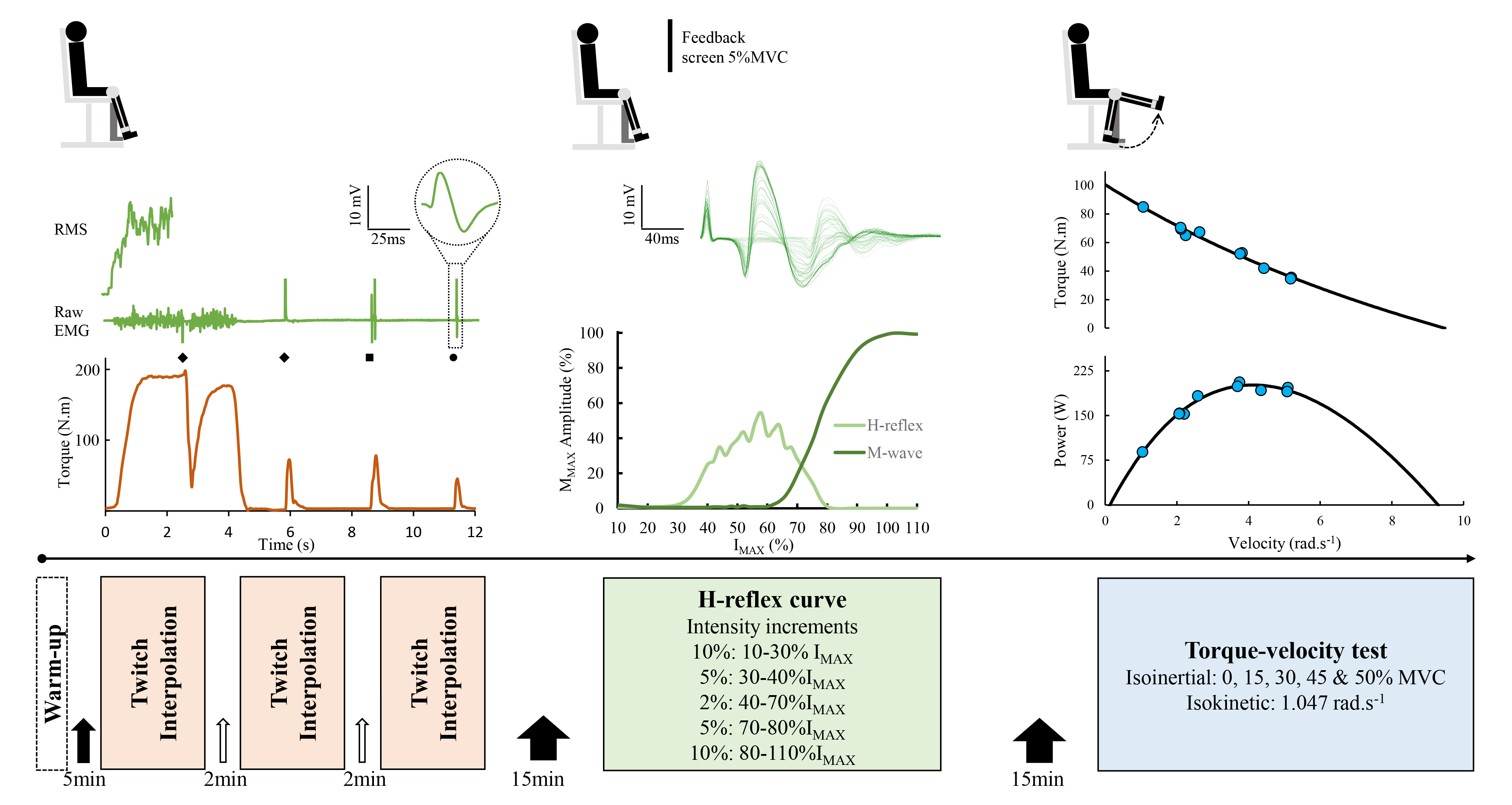
Timeline of the experimental protocol. Following the generalised warm-up, the twitch interpolation procedure was administered three times with 2 minutes rest. Torque and raw EMG of the superficial quadriceps (*vastus lateralis* example) were synchronously recorded, with RMS applied to raw EMG during MVC. High frequency 100Hz doublets (diamonds), low frequency 10Hz doublet (square) and 1Hz single pulse (circle) were applied to the femoral nerve. M-wave amplitude and duration were examined from the single pulse EMG response (inset). Following 15 minutes rest, H-reflexes were elicited in quadriceps by constructing a stimulus response curve during 5% MVC background contraction. Fifty stimulations were delivered at progressively increasing intensity, with 10-sec separating each contraction. An example raw trace of all overlaid responses is shown, with lighter lines representing responses at lower stimulation intensities. The peak-to-peak M-wave amplitude and H-reflex were plotted to construct a stimulus response curve. Following another 15 minute rest period, torque –velocity and power-velocity relationships were constructed by plotting torque and power against velocity (blue dots) measured over a series of isoinertial and one isokinetic contractions.

### 2.3 Electrical stimulation

Electrical stimulation of the femoral nerve was induced with a constant-current electrical stimulator (model DS7AH, Digitimer, Welwyn Garden City, UK) and custom-built pulse frequency generator. Voltage was set to maximum (400 V) and pulse duration to 1ms. A handheld ball-point cathode wrapped in gauze and soaked in saline (20mm diameter) and a 90mm x 50mm self-adhesive rectangular anode placed midway between the superior aspect of the iliac crest and the greater trochanter of the femur were used to stimulate the femoral nerve. At the beginning of each session, the location of the femoral nerve in the femoral triangle was determined as the position were quadriceps twitch force and M-wave responses were highest at a stimulation intensity of 50mA. This site was marked with indelible ink to ensure the stimulation site remained optimal and consistent throughout the experiment. A stimulus response curve was constructed by measuring the peak torque and peak-to-peak M-wave amplitude elicited from stimulations delivered at 10mA increments from 50mA up to the intensity which elicited no further increase in the responses (I_MAX_). Stimulation intensity for maximal evoked responses was set to 130% I_MAX_ (140 ± 28mA). For H-reflexes, stimulation intensity began at 10% I_MAX_ and was increased after every second contraction (i.e., two stimulations at each intensity) by 10% for contractions one - six, 5% between contractions six - ten, 2% between contractions ten – forty, 5% between contractions forty – forty-four, and 10% between contractions forty-four – fifty. The H-reflex protocol was slightly modified from previous protocols (44) by implementing relative increases in stimulation intensity and by asking participants to hold a 5% MVC background torque level. This approach avoided between-participant variations in the number of contractions/stimulations throughout the procedure. Moreover, piloting within a sample of our cohort demonstrated that background torque levels > 5% MVC led to fatigue development (via decrease in MVC) in some participants, which is known to impact the H-reflex response (45).

### 2.4 Surface Electromyography

Disposable pre-gelled Ag-AgCl surface electrodes (Blue sensor N, Ambu, Ballerup, Denmark) were used to record EMG signals unilaterally for *vastus lateralis* (VL), *vastus medialis* (VM) and *rectus femoris* (RF) in a bi-polar arrangement. A single 20mm Ag-AgCl ground electrode (Conmed ClearTrace ECG sensor electrode, Utica, NY) was attached to the lateral malleoli of the opposite ankle. Bi-polar surface electrodes were positioned at an inter-electrode distance of 20mm and aligned parallel to the muscle fibres in accordance with the recommendations of the SENIAM project (46). Prior to placement of electrodes, the skin was prepared by shaving, abrading and cleaning with an alcohol swab. EMG electrodes were secured with adhesive tape to ensure good contact with the skin and to reduce movement artefact. Indelible ink was used to mark electrode location at the end of the first session and participants were asked to avoid cleaning the area before the second session, ensuring that electrode location was consistent between sessions.

### 2.5 Data acquisition and analysis

All torque (N.m), angle (rad), velocity (rad.s^-1^), EMG (mV) and stimulation pulse analog data were synchronously acquired at 4000Hz and digitally converted with a Powerlab 8/30 (ADinstruments, Australia) in conjunction with a PC running Labchart software (version 8.1.24, ADinstruments, Australia). Raw EMG signals were amplified (gain: 1000, BioAmp ADinstruments, Australia) and processed with a bandpass filter (fourth order Butterworth 20 – 500Hz) and 50ms root mean square (RMS) for data recorded during MVC. All offline processing was completed using Spike2 software (version 7.13, Cambridge Electronic Design, Cambridge, UK).

#### 2.5.1 Isometric voluntary and evoked torque responses

Gravity correction of the recorded isometric torque data was applied by adding the torque generated by the passive weight of the leg and dynamometer attachment at 75° knee flexion. During MVC, average torque (N.m) and RMS (mV) were calculated over a 500ms time window preceding superimposed stimulation, and superimposed twitch torque (SIT) was calculated as the change in torque within an ∼120ms time window following stimulation. From the potentiated resting twitches at 100Hz, 10Hz and 1Hz, peak torque (Pt), maximal rate of torque development (MRTD – maximum of the torque-time slope following 100Hz stimulation (1ms time constant)) and the ratio between Pt_10Hz_ and Pt_100 Hz_ (Pt_10:100_) (47) was calculated. The equation 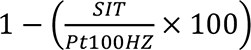 was used to calculate voluntary activation (%) (33). The average value from all three assessments were used for analysis unless MVC force was < 90% of the highest value recorded in a particular session (in which case it was discarded).

#### 2.5.2 Evoked electromyography responses

From the maximal M-wave (MMAX) responses obtained from the raw EMG during 1Hz resting twitches, peak-to-peak amplitude and peak-to-peak duration were calculated for all quadriceps muscles. RMS during MVC was normalised to MMAX (RMS.MMAX %) to remove the contribution of peripheral factors on the RMS signal and to permit better representation of overall neural drive (32, 48). From the H-reflex procedure, peak-to-peak amplitude of the H-reflex and M-wave were calculated from each stimulation given to construct a stimulus response curve (**Figure 1**). H_MAX_ was determined as the highest peak-to-peak amplitude relative to maximal M-wave amplitude (H_MAX_.M_MAX_) (44).

#### 2.5.3 Torque-velocity and power-velocity relationships

Concentric torque data was gravity offset by multiplying the passive torque generated by the weight of the leg at 10° flexion (0 = full extension) by the cosine of the recorded knee angle (49). An additional power channel (W) was calculated by multiplying torque (N.m) by velocity (rad.s^-1^). For all maximal efforts, average torque, velocity and power were extracted from torque onset (3x standard deviations above baseline values) to knee angle of 30° flexion. This angle was chosen to avoid increased resistance generated by the braking torque (i.e. cushioning) of the dynamometer toward full knee extension during high-velocity contractions. The average value was chosen over the peak due better reliability in the outcomes and closer associations to physical function (26). Torque-velocity (T-V) relationships were modelled by re-writing the original hyperbolic equation developed by Hill (50) (non-linear least squares method):

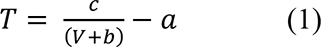

where T is the torque produced at a given velocity V, and a, b and c are constants. From the T-V relationship the following variables were extracted: 1) maximal torque (T0) at the y-axis intercept when V = 0, 2) maximal velocity (V0) at the x-axis intercept calculated as 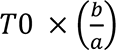, 3) optimal torque (T_opt_) at maximal power calculated as (*a*^2^ + (*a* × *T*0))^0.5^ − *a*, and 4) optimal velocity (V_opt_) at maximal power calculated as 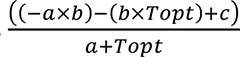 (1) (51-53). Power-velocity relationships were also modelled using the equation:

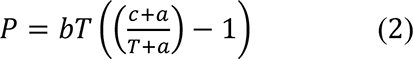

where P is power produced at a given torque and velocity T and a, b and c are constants. Maximal power (P_MAX_) occurred at the apex of the power-velocity relationship and was calculated as 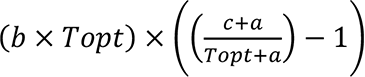 (54). The single isokinetic effort at 1.047 rad/s (i.e., 60 deg/s) was included in the modelling procedure to increase the validity of torque estimates at low contraction velocities. This could not be achieved with an isoinertial load beyond 50% MVC, as there was large inter-individual variability in the capacity for participants to achieve the required ROM due to the limiting effects of the length-tension relationship (55). Both efforts completed at each load/velocity were included in modelling procedures unless they deviated by >5%, in which case, the highest value was included.

### 2.6 Statistics

All statistical analysis was performed using R Software (v4.2.2; R Core Team 2022). Normal distribution of the original data and delta between sessions was assessed with histograms, Q-Q plots, and Shapiro-Wilks test, with natural log transformations applied if data displayed a non-normal distribution. Homoscedasticity of all data was confirmed with the Breusch-Pagan test. Generalized linear mixed models were used to assess differences in the outcomes between sessions, with participant set as the random effect. An additional fixed effect of muscle was included for EMG outcomes to determine differences in the delta between sessions between *vastus lateralis, vastus medialis* and *rectus femoris* muscles (i.e. session x muscle interaction). The relationship between delta in the outcomes between session one and two with age of the participants was assessed using linear regression. Significance was set at P < 0.05 for all statistical tests.

Relative reliability (i.e. participant position within the sample) between sessions one and two was assessed by the intraclass correlation co-efficient (ICC) with a two-way random effects model for single-measure reliability (56). ICC values between 0.8 – 1 were considered ‘excellent’, 0.6 – 0.8 as ‘good’ and < 0.6 as ‘poor’ (57). Absolute reliability (i.e., variability in participant scores) between session one and two was assessed by calculating the typical error, coefficient of variation, and the mean difference with 95% limits of agreement. Typical error (TE) in the original unit of measurement was calculated by dividing the SD of the absolute difference between sessions one and two by 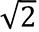, and the coefficient of variation (CoV) in percent was calculated with the formula (*SD*/*Mean*) × 100. Bland-Altman plots were constructed to determine the systematic error (i.e. bias toward higher or lower outcomes during session one or two) and random error (i.e. inherent biological or mechanical variations), by plotting the individual participant difference in the outcome between session one and two against the mean, and the 95% lower and upper limits of agreement (LOA), respectively (58, 59). If log transformation of original data were required, the antilog of the mean difference and LOA was used to transform log values to a percentage difference (58, 60). LOA ratios (%) were calculated to allow direct comparison with previous results when necessary (58).

## 3. Results

### 3.1 Isometric voluntary and evoked forces

Isometric voluntary and evoked force measurements were similar between the sessions (all P > 0.05). All outcomes displayed ICCs > 0.8 (**Table 1**). CoVs were 2.6% for VA and < 5.5% for peak twitch forces and MRFD. No notable systematic bias was observed for any of the voluntary or evoked force outcomes (**Figure 2**).

**Figure 2.**
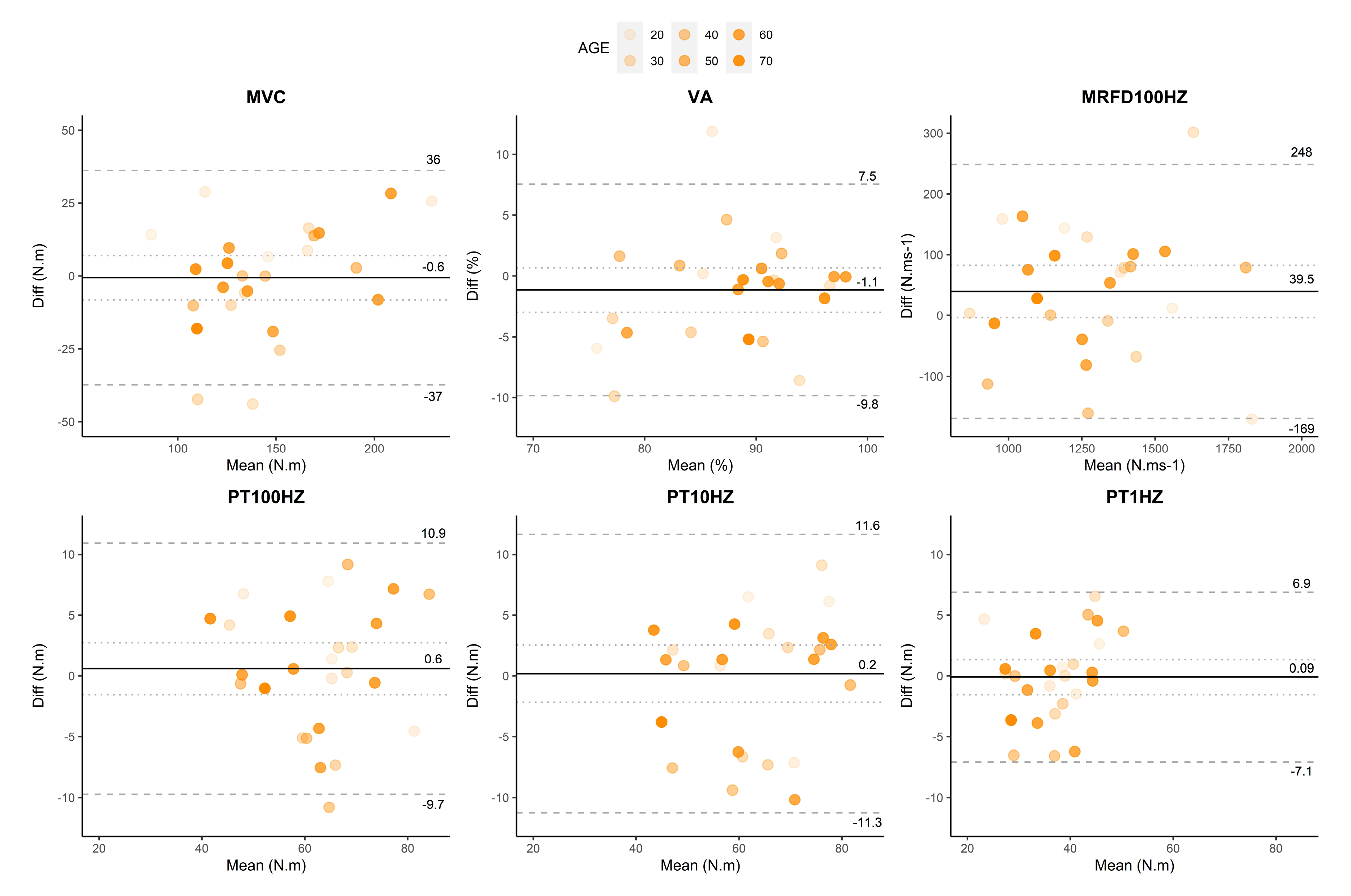
Bland Altman plots with 95% level of agreement for voluntary and evoked forces. The mean value for session one and two is plotted on the x axis, whereas the session one minus session two difference (Diff) is plotted on the y axis. The solid line represents the grand mean difference for all participants and the dotted line shows the 95% confidence interval. The dashed lines represent the lower and upper limits of agreement in original unit of measurement. Shading of the individual measures increases as a function of age.

**Table 1.** Mean ± standard deviation, inter-session reliability, and regression (session delta vs. age) for all outcome

|  | <u>Session #1</u> | <u>Session #2</u> | <u>Inter-session reliability</u> |  |  | <u>Age effect</u> |
| --- | --- | --- | --- | --- | --- | --- |
|  | Mean (SD) | Mean (SD) | Typical error | CV (%) | ICC (lower, upper 95% CI) | R<sup>2</sup> |
| **Isometric voluntary and evoked forces** |  |  |  |  |  |  |
| MVC (N.m) | 144.85 (39.05) | 146.77 (32.74) | 13.26 | 7.24 | 0.869 (0.728, 0.939) | 0.002 |
| SIT (N.m) | 7.92 (4.83) | 7.36 (4.12) | 1.78 | 18.35 | 0.842 (0.677, 0.927) | <0.00 |
| VA (%) | 87.44 (7.49) | 88.48 (6.48) | 3.13 | 2.60 | - | <0.00 |
| Pt<sub>100Hz</sub> (Nm) | 63.04 (11.27) | 62.53 (11.06) | 3.72 | 4.79 | 0.893 (0.777, 0.951) | <0.00 |
| Pt<sub>10Hz</sub> (Nm) | 62.3 (12.69) | 62.5 (12.49) | 4.13 | 5.62 | 0.897 (0.783, 0.952) | 0.03 |
| Pt<sub>1Hz</sub> (Nm) | 37.11 (7.67) | 37.27 (6.54) | 2.51 | 5.26 | 0.881 (0.752, 0.945) | 0.02 |
| Pt<sub>100Hz:10Hz</sub> Ratio | 1.02 (0.11) | 1.01 (0.09) | 0.04 | 3.12 | 0.847 (0.69, 0.928) | 0.11 |
| MRFD<sub>100Hz</sub> (Nm.s<sup>-1</sup>) | 1311.5 (2260.5) | 1279.3 (255.5) | 75.40 | 4.92 | 0.908 (0.799, 0.958) | <0.00 |
| **Isometric voluntary and evoked EMG** |  |  |  |  |  |  |
| *Vastus lateralis* |  |  |  |  |  |  |
| RMS (mV) | 0.217 (0.119) | 0.220 (0.108) | 0.02 | 9.22 | 0.95 (0.89, 0.978) | 0.02 |
| RMS.M<sub>MAX</sub> (%) | 5.08 (2.61) | 5.24 (2.60) | 2.09 | 10.97 | 0.846 (0.667, 0.933) | 0.06 |
| H<sub>MAX</sub>.M<sub>MAX</sub> (%) | 23.22 (14.14) | 23.02 (16.50) | 3.69 | 11.83 | 0.966 (0.914, 0.987) | <0.00 |
| M<sub>MAX</sub> (mV) | 4.52 (2.34) | 4.62 (2.23) | 0.52 | 8.34 | 0.945 (0.872, 0.977) | 0.01 |
| M<sub>MAX</sub> Duration (ms) | 9.87 (2.9) | 10.03 (2.75) | 1.0 | 4.09 | 0.963 (0.915, 0.985) | <0.00 |
| *Vastus medialis* |  |  |  |  |  |  |
| RMS (mV) | 0.35 (0.22) | 0.36 (0.20) | 0.03 | 7.89 | 0.966 (0.927, 0.985) | 0.01 |
| RMS.M<sub>MAX</sub> (%) | 3.45 (1.43) | 3.7 (1.20) | 0.65 | 14.39 | 0.758 (0.528, 0.885) | <0.00 |
| H<sub>MAX</sub>.M<sub>MAX</sub> (%) | 19.51 (13.49) | 22.59 (13.71) | 4.88 | 22.39 | 0.857 (0.656, 0.945) | 0.06 |
| M<sub>MAX</sub> (mV) | 9.57 (3.27) | 9.58 (3.48) | 1.36 | 12.43 | 0.792 (0.585, 0.903) | 0.01 |
| M<sub>MAX</sub> Duration (ms) | 10.07 (2.31) | 10.72 (2.88) | 1.0 | 6.61 | 0.877 (0.741, 0.944) | 0.06 |
| *Rectus femoris* |  |  |  |  |  |  |
| RMS (mV) | 0.18 (0.82) | 0.18 (0.81) | 0.02 | 13.76 | 0.852 (0.698, 0.931) | 0.03 |
| RMS.M<sub>MAX</sub> (%) | 6.16 (1.97) | 6.16 (1.76) | 0.96 | 13.89 | 0.721 (0.38, 0.89) | 0.05 |
| H<sub>MAX</sub>.M<sub>MAX</sub> (%) | 18.80 (9.94) | 15.65 (7.23) | 5.66 | 16.73 | 0.644 (0.151, 0.887) | 0.06 |
| M<sub>MAX</sub> (mV) | 2.97 (0.75) | 2.73 (1.09) | 0.47 | 15.04 | 0.561 (0.139, 0.813) | 0.29* |
| M<sub>MAX</sub> Duration (ms) | 9.95 (2.42) | 10.02 (1.87) | 1.0 | 6.01 | 0.899 (0.734, 0.963) | 0.01 |
| **F-V and P-V relationships** |  |  |  |  |  |  |
| PMAX (W) | 179.7 (42.16) | 183.33 (42.16) | 9.80 | 4.03 | 0.945 (0.881, 0.975) | <0.00 |
| T0 (Nm) | 119.78 (27.73) | 115.84 (25.47) | 6.55 | 4.82 | 0.938 (0.865, 0.972) | <0.00 |
| Topt (Nm) | 48.40 (9.13) | 47.83 (8.52) | 2.43 | 3.75 | 0.928 (0.843, 0.967) | <0.00 |
| V0 (rad.s<sup>-1</sup>) | 8.50 (1.88) | 8.63 (1.92) | 0.62 | 5.97 | 0.897 (0.781, 0.953) | <0.00 |
| Vopt (rad.s<sup>-1</sup>) | 3.42 (0.46) | 3.54 (0.48) | 0.17 | 4.13 | 0.852 (0.692, 0.932) | <0.00 |

measures.

### 3.2 Voluntary EMG, M-wave and H-reflex

Original data from eight of the fifteen EMG variables displayed a non-normal distribution. Consequently, all EMG variables were log transformed prior to analysis to permit investigation into intermuscular differences in the outcomes and to permit meaningful interpretation of the LOA (61). EMG, M-wave, and H-reflex responses were similar for session one and two for all quadriceps muscles (all P > 0.05) and the difference in the outcomes between sessions was not significantly different between the muscles (i.e., no interaction effects; all P > 0.05). ICCs for all outcomes measured in *vastus lateralis* muscle were > 0.8 (**Table 1**). RMS, H_MAX.MAX_ and M_MAX_ duration measured in *vastus medialis* and RMS and M_MAX_ duration measured in *rectus femoris* also showed ICCs > 0.8. CoVs were lower in *vastus lateralis* compared to *vastus medialis* and *rectus femoris* muscle for 4/5 EMG outcomes, with all *vastus lateralis* outcomes showing a CoV of < 12%. *Vastus lateralis* showed no notable systematic bias (all < ∼1%) and less random error compared to *vastus medialis* (with exception of RMS) and *rectus femoris* muscles for all outcomes (**Figure 3**). Feasibility for M-wave/H-reflex responses was 81/73% for *vastus lateralis*, 96/65% for *vastus medialis*, and 65/42% for *rectus femoris*.

**Figure 3.**
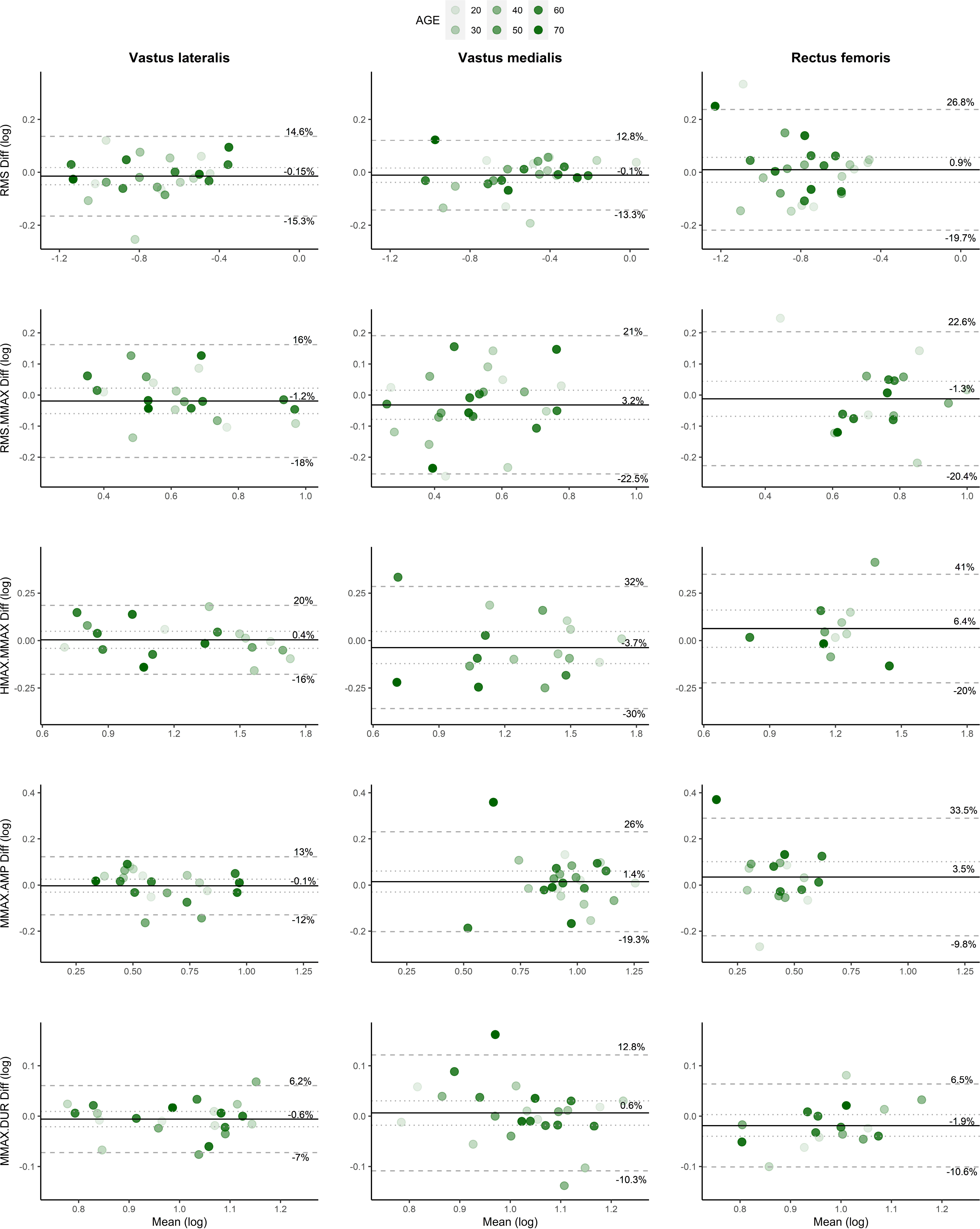
Bland Altman plots with 95% level of agreement for *vastus lateralis* (left column) *vastus medialis* (centre column) and *rectus femoris* (right column) muscles. The mean value for session one and two is plotted on the x axis, whereas the difference between session one and two is plotted on the y axis. The solid line represents the grand mean difference for all participants and the dotted line shows the 95% confidence interval. The dashed lines represent the lower and upper limits of agreement as a % difference (antilog of log transformed data). Shading of the individual measures increases as a function of age.

### 3.3 Torque-velocity and power-velocity relationships

All outcomes extracted from the torque-velocity and power-velocity relationships were similar between sessions (all P > 0.05). ICCs were > 0.8 for all outcomes (**Table 1**). CoVs for V0, T0 and P_MAX_ were ∼6%, ∼5% and ∼4%, respectively. No notable systematic bias was identified for any of the outcomes (**Figure 4**).

**Figure 4.**
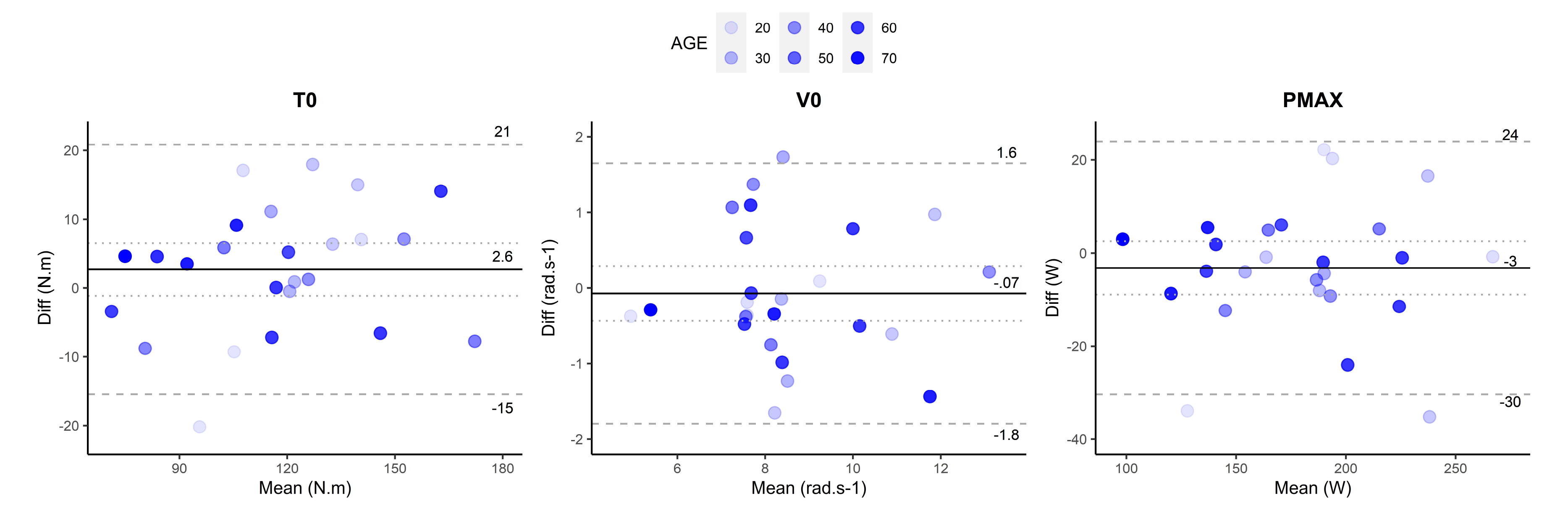
Bland Altman plots with 95% level of agreement for variables extracted from the torque-velocity and power-velocity relationships. The mean value for session one and two is plotted on the x axis, whereas the difference between session one and two is plotted on the y axis. The solid line represents the grand mean difference for all participants and the dotted line shows the 95% confidence interval. The dashed lines represent the lower and upper limits of agreement in original unit of measurement. Shading of the individual measures increases as a function of age

## 4. Discussion

All mechanical outcomes extracted from isometric maximal voluntary and evoked contractions demonstrated excellent relative and absolute reliability between session one and two. Similarly, all mechanical outcomes derived from the force-velocity and power-velocity relationships constructed from maximal concentric contractions demonstrated excellent reliability between the sessions. *Vastus lateralis* muscle exhibited higher ICCs (all > 0.8) and lower CoVs (all < 12%) compared to *vastus medialis* and *rectus femoris* muscles. Age was not correlated with the mean difference between sessions for any of the outcomes.

### 4.1 Isometric voluntary and evoked forces

#### 4.1.1 MVC

Quadriceps MVC, as a gold-standard measure of maximal muscle strength, demonstrated excellent inter-session reliability with an ICC of 0.869 and CoV of 7.2%. Random error in strength measurements may be exacerbated in older adults due to large between-subject variability in neuromuscular performance (34) or as they are generally less accustomed to high-force contractions compared to younger adults (22, 62). Our findings indicate that random error in MVC measurements may be dependent on overall voluntary strength capacity rather than age, as the mean difference in MVC between the sessions seemed higher when the mean MVC was lower, with no association with age (**Figure 2**). Moreover, the overall mean difference at force levels < ∼150 N.m appeared in favor of higher values during the second session, indicating that females with lower strength capabilities may benefit from a second session to record maximal quadriceps isometric voluntary strength. Nonetheless, systematic bias was absent from MVC measurements in accordance with previous observations from isometric contractions (22). Our method of measuring quadriceps MVC prior to impending supramaximal stimulation of the peripheral nerve did not appear to increase the random error in the measurement, as reliability was similar to previous findings from older females and when stimulation of the nerve was not applied (12, 13). Thus, the adopted method is advantageous in providing an elaborate evaluation of neuromuscular function without compromising MVC reliability. The reported LOA are acceptable for quantifying age-related differences in quadriceps MVC (63, 64).

#### 4.1.2 Voluntary Activation

ICCs for quadriceps VA were not calculated due to the ceiling effect of the measurement (15, 65), however, the CoV was 2.6%, and the upper and lower LOA were 7.5% and −9.8% (LOA ratio = 9.8%). Systematic bias was small, with a mean difference of 1.1% toward higher VA measurements during the second session (22). These results represent slightly higher inter-session reliability and less random error in quadriceps VA measurements compared to younger males and females (15-17) and older patients with knee osteoarthritis (40). However, many methodological factors can influence VA measurements (22, 66), which makes a physiological explanation for the small discrepancy in reliability difficult. The reported LOA must be considered when evaluating age-related deficits in voluntary activation of the quadriceps, as significant reductions of little as ∼5% (67, 68) up to ∼11% (69) have previously been reported.

#### 4.1.3 Resting twitch responses

All responses elicited from single and double pulse (100Hz and 10Hz) stimulations exhibited excellent reliability, evidenced by ICCs ranging from ∼0.85 – 0.91 and CoVs of < ∼5%. For the peak force responses, there was no evidence of systematic bias (≤ 0.6 N.m) and random error was < ∼11 N.m. These results are either similar or more reliable than previous findings obtained from younger males and females (15, 16), and comparable to older adults with knee osteoarthritis (40). Age may reduce random error in twitch responses as older adults have less muscle potentiation following a pre-conditioning contraction (70), which is dependent on type II fibres that preferentially atrophy into older age (34) and highly variable (71). Although single pulse twitches were anecdotally more tolerable than doublets, single pulse twitch force was ∼2/3rds the magnitude of the doublet peak force and thus did not represent the true potential of intrinsic muscle contractility. High and low frequency twitch forces are also more valuable than single twitch force responses as age-related deficits in factors such as neuromuscular propagation (e.g. at the neuromuscular junction and along the sarcolemma; 100Hz) and excitation-contraction coupling (e.g. myofibrillar calcium kinetics and sensitivity; 10HZ) can be evaluated (32, 34, 72). It is also worth noting no participants withdrew their voluntary consent to participate in the electrical stimulation protocol and all returned for the second re-test session. Thus, the stimulating procedure was feasible in evaluating peripheral contractile skeletal muscle characteristics in healthy younger and older females. However, for more frail elderly or clinical groups, alternate (albeit limited) methods may be more appropriate (e.g. reducing supramaximal intensity) (32, 40).

### 4.2 Electromyography

All quadriceps muscles showed excellent reliability for RMS, though *vastus lateralis* was the only muscle to show excellent reliability for RMS.M_MAX_ (ICC = 0.85). M-wave responses and H_MAX_.M_MAX_ also showed excellent reliability in *vastus lateralis* (ICCs > 0.95) and CoVs for were lower compared to the other superficial quadriceps. These outcomes were more reliable than reported previously for younger males (18, 44, 61, 73) and M_MAX_ was comparable to older males and females with knee osteoarthritis (40). The 5% MVC background force level adopted during H-reflex procedures may have improved reliability relative to previous studies, as females recruit more stable low threshold motor units at higher firing rates to generate submaximal force levels compared to males (5). Thus, it may be more reliable to elicit quadriceps H-reflex responses at low background force levels when assessing females. The LOA of agreement are acceptable for detecting age-related changes in RMS and M-wave for both *vastii* muscles but may not be sensitive enough for *vastus medialis* H-reflex amplitude (68). Although age-related effects on *vastus lateralis* H-reflex amplitude have not previously been reported, H-reflex amplitude for the soleus ankle plantar-flexor muscle decreases as a function of age with deficits of ∼20% reported in older adults (74). It is important to note that direct comparison on the reliability of voluntary and evoked EMG outcomes between studies is troublesome, as reliability is influenced by a number of factors including muscle length (37, 61, 75) and electrode location (61). Thus, the reliability of voluntary and evoked EMG responses should be established for specific protocols prior to experimentation.

*Vastus lateralis* EMG appeared more reliable compared to *vastus medialis* and *rectus femoris* muscles. ICCs were higher, CoVs were lower, and the random error was smaller in *vastus lateralis* for 4/5 of the outcomes (**Table 1** and **Figure 3**), despite not reaching statistical significance. *Vastus lateralis* outcomes were also more reliable than findings reported for younger healthy males (15, 18, 44, 61, 76). Thus, when evaluating age-related differences in female neuromuscular function, voluntary and evoked EMG outcomes may be more reliable in *vastus lateralis*. Of all the superficial quadriceps, *vastus lateralis* possesses a higher cross sectional area (irrespective of age) (41, 77), is the greatest contributor to the knee extensor moment (78) and the most reliable predictor of force output during knee extension and leg extension movements (79). Moreover, although *vastus medialis* shares some common neural drive with *vastus lateralis* (and theoretically should exhibit similar levels of reliability) this is highly variable (80, 81) and potentially exacerbated in females (42). The poor reliability and feasibility for *rectus femoris* evoked EMG responses was somewhat expected (44) possibly due to their bipennate architecture and close proximity to the site of stimulation.

### 4.3 Torque-velocity and power-velocity relationships

All quadriceps torque-velocity and power-velocity characteristics exhibited excellent inter-session reliability (ICCs > 0.8). Inter-session reliability of quadriceps T0 between the two sessions was similar to measurements of quadriceps 1RM (11) (although in that study, significant differences between the two sessions was observed) and isokinetic torques generated at 60°/s and 180°/s (10) in young and elderly males and females. Moreover, V0 and P_MAX_ showed lower CoVs then when measured separately from single isoinertial loads corresponding to 20%, 25% or 50% MVC in older males and females (82, 83). Our results show that measures of quadriceps maximal torque, velocity and power obtained from a torque-velocity test may provide more reliable and elaborate information regarding neuromuscular function compared to 1RM or isokinetic tests. Indeed, functional capacity in older adults is more strongly related to maximal muscular power than muscle strength (23), and shortening velocity is a primary determinant of muscular power (23, 24).

### 4.4 Considerations

Although it is unclear if a third testing session could further improve the reliability of the outcomes, previous studies in younger males have shown negligible changes in maximal voluntary and evoked forces (17-19) and force-velocity outcomes (84). Thus, inconveniences imposed on participants (e.g. time commitment and exposure to peripheral nerve stimulation techniques) can be minimized, which increases the feasibility of implementing these techniques into future research protocols.

It is also worth noting that the average duration between testing sessions of 9 ± 7 days may have in some cases led to eumenorrheic females (n = 7) being assessed at different phases of the menstrual cycle. Menstrual cycle phase was not controlled for as this was not expected to influence the mechanical outcomes (22). The menstrual cycle phase may however influence motor unit discharge rate (85) and be a source of random error in EMG outcomes, but more research is needed using advanced neurophysiological techniques (e.g. high-density EMG) (6). Testing all eumenorrheic females in the same phase of their menstrual cycle would however have increased the test-re-test duration to the duration of one menstrual cycle (i.e. ∼28 days), thus introducing further and potentially larger sources of variability.

## 5. Conclusions

This is the first study to provide a comprehensive analysis of the test-retest reliability of quadriceps maximal voluntary and evoked forces and electromyography outcomes used to evaluate the aging neuromuscular system and adaption to exercise, with special reference to females. The results demonstrate that reliable and accurate measurements can be obtained in one testing session with no effect of age. This is important for future research aiming to quantify age-related degeneration in neuromuscular function and/or neuromuscular adaptation to exercise at different stages of the female lifespan.

## Acknowledgements

The authors would like to thank Ms. Brianna Gatto for her role in managing participant recruitment and co-ordination.

## Author contributions

SOB, DH and SL conceived and designed research, SOB performed experiments, SOB analysed data, SOB interpreted results of experiments, SOB prepared figures, SOB drafted manuscript, SOB and SL edited and revised manuscript, SOB, DH and SL approved final version of manuscript.

---

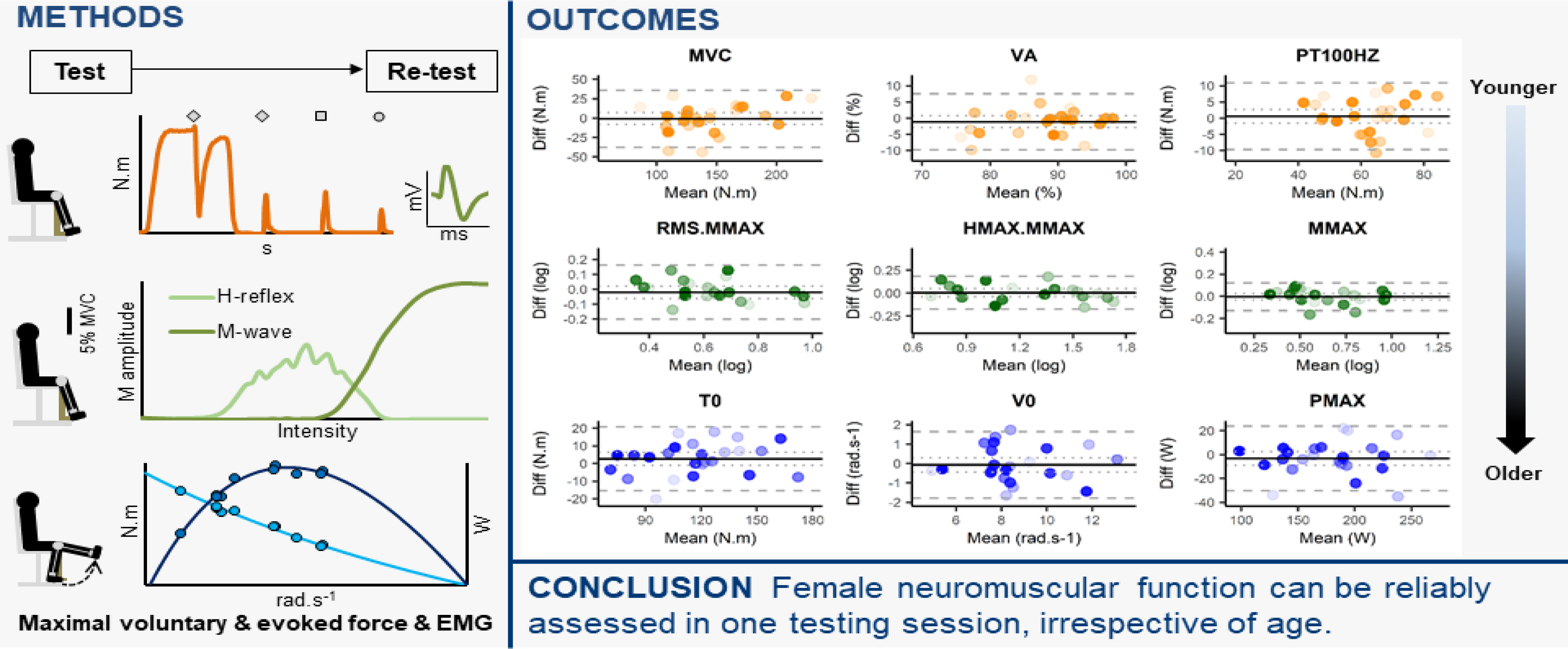

